# Expanding the coverage of regulons from high-confidence prior knowledge for accurate estimation of transcription factor activities

**DOI:** 10.1101/2023.03.30.534849

**Authors:** Sophia Müller-Dott, Eirini Tsirvouli, Miguel Vázquez, Ricardo O. Ramirez Flores, Pau Badia-i-Mompel, Robin Fallegger, Astrid Lægreid, Julio Saez-Rodriguez

## Abstract

Gene regulation plays a critical role in the cellular processes that underlie human health and disease. The regulatory relationship between transcription factors (TFs), key regulators of gene expression, and their target genes, the so called TF regulons, can be coupled with computational algorithms to estimate the activity of TFs. However, to interpret these findings accurately, regulons of high reliability and coverage are needed. In this study, we present and evaluate a collection of regulons created using the CollecTRI meta-resource containing signed TF-gene interactions for 1,183 TFs. In this context, we introduce a workflow to integrate information from multiple resources and assign the sign of regulation to TF-gene interactions that could be applied to other comprehensive knowledge bases. We find that the signed CollecTRI-derived regulons outperform other public collections of regulatory interactions in accurately inferring changes in TF activities in perturbation experiments. Furthermore, we showcase the value of the regulons by investigating hallmarks of TF activity profiles inferred from the transcriptomes of three different cancer types. Overall, the CollecTRI-derived TF regulons enable the accurate and comprehensive estimation of TF activities and thereby help to interpret transcriptomics data.

**GRAPHICAL ABSTRACT:** 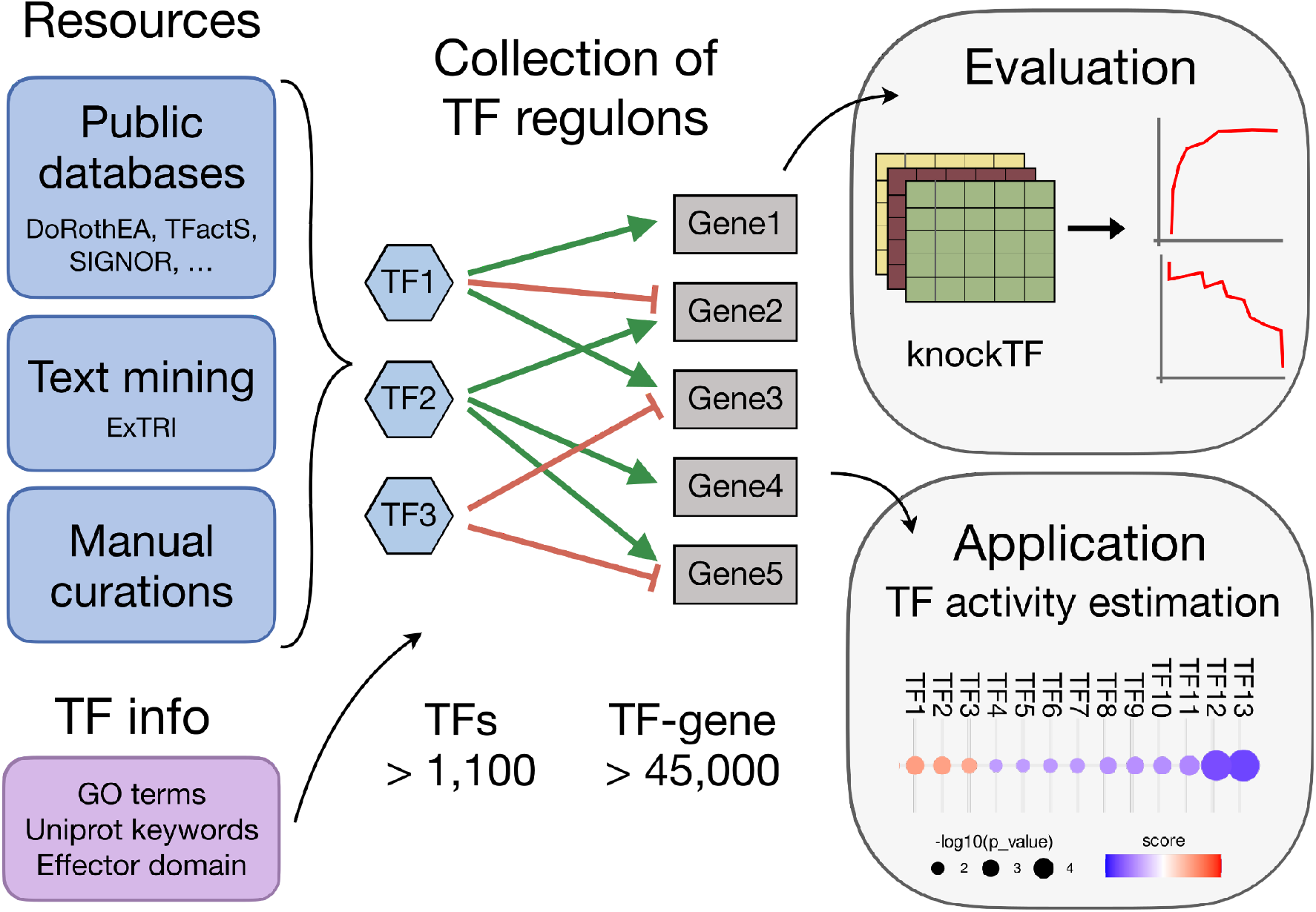

## INTRODUCTION

The regulation of gene transcription plays a fundamental role in development, cell differentiation, tissue homeostasis, and various physiological processes and is crucial for the coordinated response of cells to both internal and external signals. As such, its dysregulation can contribute to the development of numerous diseases, including cancer, autoimmunity, neurological disorders, developmental syndromes, diabetes, or cardiovascular disease (1). In particular, deregulated activity of transcription factors (TFs) - key regulators of transcription - has been implicated in the development of cancer and can generally alter the core autoregulatory circuitry of a cell (1–3). Transcription factors bind to specific regions of the DNA and together with cofactors and other proteins influence the transcriptional rate of a specific set of target genes (TGs) collectively known as the TF’s regulon (4). The combined interactions of all TFs to their target genes are referred to as a gene regulatory network (GRN), a simplified representation of the underlying regulatory circuits (5). Coupling GRNs with activity inference algorithms (6) can facilitate the interpretation of transcriptomics data and provide a more effective means of understanding the underlying regulatory mechanisms in the system of interest. Among other, TF activity estimation has been used to better understand aging (7) and to relate TF activities to treatment efficacies and resistance mechanisms (8, 9), to patient survival (10) and to cancer morphological features (11). Since the choice of TF-regulons can substantially affect the results (12), it is important to use TF-regulons of high quality, minimizing false-positive interactions, while having the highest coverage possible to not miss potentially relevant TFs.

Various methods are available for identifying TF regulons, both on a small and large scale. Experimentally, high-resolution identification of TF binding sites *in vivo* can be achieved using chromatin immunoprecipitation followed by high-throughput DNA sequencing (ChIP-seq) (13) and DNase I hypersensitivity coupled with DNA sequencing (DNase-seq) (14). Despite identifying TF-DNA binding events in their native environment, binding events might not correspond to actual changes in the expression of the target gene, and do not take into account other cofactors that bind indirectly to the target genes (15). *In silico*, the prediction of TF-gene interactions can be done using the genomic sequence recognised by each TF, also known as binding motifs (16, 17). Such methods involve probing the entire genome for regions that contain these binding motifs to identify potential target genes. However, this approach is limited to TFs with known binding motifs and does not account for context-specific interactions, where regulatory interactions take place only in a specific cell type or condition. Furthermore, GRNs can be inferred in a data-driven manner, as in co-expression analysis where the correlation between the expression patterns of a TF and its potential target genes is investigated (18). Lastly, manual curation of TF-gene interaction from the literature is another common strategy. Such curation efforts are usually incorporated in databases such as IntAct (19), SIGNOR (20), and TRRUST (21). While being very attractive for their high quality, manual curations are hard to come by since curation is a generally cumbersome task, and curated databases rarely overlap due to the different curation standards and protocols (12, 22). For that, curation efforts can be significantly enhanced with the aid of text-mining for the identification of TF-gene interactions, as previously shown (23).

Despite the availability of the afore-described methods, the lack of a general consensus on the inference of TF-gene interactions remains a challenge as each approach has its own strengths and limitations. A few frameworks to create TF regulons based on a combination of resources have been proposed. Such frameworks include DoRothEA (12), which combines TF-gene interactions identified by ChIP-seq experiments, inferred interactions by gene expression and TF binding motifs and manual curation, and ChEA3 (24), which primarily contains co-expression and ChIP-seq-inferred interactions. Other examples include Pathway Commons, which is a resource that integrates various types of interactions (i.e. biochemical, complex binding and physical interactions between proteins, RNA, DNA and small molecules) (25) and RegNetwork, which compiles experimentally observed or motif-based predicted interactions among TFs, microRNAs and target genes (26). However, such meta-resources can include a high number of false interactions due to the use of high false-positive generating methods (i.e. co-expression and co-occurrence) (27), and, with the exception of DoRothEA, they do not include the information about the sign of interactions (i.e. activating or inhibiting interactions).

In this study, we introduce a set of TF regulons created using information on TF-gene interactions from the CollecTRI (Collection of Transcription Regulation Interactions) meta-resource (23) which integrates multiple sources of interaction data, including public databases, text mining, and manual curation. The CollecTRI-derived regulons represent 45,856 signed TF-gene interactions for 1,183 TFs. Additionally, we propose a workflow for defining the sign of interactions (activating or repressing) based on i) information about the sign curated in the resources compiled in CollecTRI, on ii) prior knowledge about the TFs activating and/or repressing roles and on iii) regulon properties which can also be applied to other comprehensive knowledge bases. We benchmarked the performance of CollecTRI-derived regulons against TF regulons from four other meta-resources: DoRothEA, ChEA3, RegNetwork and Pathway Commons. The CollecTRI-derived regulons outperformed the other networks by accurately inferring changes in TF activities in TF perturbation experiments collected in the KnockTF data (28). Lastly, we demonstrated the value of the CollecTRI-derived regulons through a case study, where we estimated the differential activities of TFs in three cancer types using public data from the Clinical Proteomic Tumor Analysis Consortium (CPTAC). We were able to estimatethe differential activities of TFs with known roles in cancer biology and the studied cancer types, highlighting the value of the extended coverage of CollecTRI-derived regulons. In summary, we provide a new high-confidence, high-coverage collection of TF-regulons that we make freely available via OmniPath (https://omnipathdb.org/) (29) and the DoRothEA package (https://saezlab.github.io/dorothea/) (12).

## MATERIAL AND METHODS

### TF-gene data sources

The CollecTRI source data were introduced in (23) and have since been updated by gathering more recent data from SIGNOR and GO, and by adding three new resources: DoRothEA_A, Pavlidis 2021, and the NTNU Curated subset of ExTRI. Pavlidis consists of information extracted from the supplemental materials of the publication. The code that implements the gathering and merging of the data is available within the ExTRI Rbbt workflow (https://github.com/Rbbt-Workflows/ExTRI). Each data source is processed to a common format with transcription factors and target genes expressed as human gene symbols. For databases that list genes and proteins of different organisms the UniProt protein to protein identifier equivalences from the Protein Information Resource (https://proteininformationresource.org/) were used to identify the proper human protein, which was then translated to its gene symbol. Each resource was processed to update all gene symbols to the most recent version (gene set in Ensembl release 109 from February 2023).

When merging databases, entries for AP1 and NFKB complex members are allowed to match the complex names so that their information is merged across them. The result of this process is a large table where each TRI (TF-gene pair) is listed with the databases in which it is present, along with information from those databases, such as mode of regulation, when available, and the PubMed ID (PMID) from which the TRI was curated/extracted. Each database thus provides a list of PMIDs as supporting evidence for each TRI.

### Filtering TF-gene interactions

CollecTRI, as well as all other GRNs included, were filtered to contain only transcription factor (TF)-gene interactions from TFs classified as DNA-binding (dbTFs), co-regulatory (coTFs) or general initiation (GTFs). dbTFs were downloaded from Lambert et al.

(https://ars.els-cdn.com/content/image/1-s2.0-S0092867418301065-mmc2.xlsx), Lovering et al. (https://ars.els-cdn.com/content/image/1-s2.0-S1874939921000833-mmc1.xlsx) and TFclass (http://tfclass.bioinf.med.uni-goettingen.de/suppl/tfclass.ttl.gz). For coTFs and GTFs, we retained all proteins annotated in Gene Ontology (GO) (30) with the term GO:0003712 or GO:0140223 or any of their daughter terms, respectively, through QuickGO (31).

### Assigning the TF-gene mode of regulation

For each TF-gene interaction from the CollecTRI meta-resource the PMIDs were aggregated across databases. Each PMIDs is considered evidence of whichever mode of regulation is specified in that database, if any. Database entries that are not supported by PMIDs are thus not considered when building the TF regulons. PMIDs can only count as evidence for a TF-gene interaction once, even if they were used to source entries in several databases. Only in the infrequent case of the same PMID featured as supporting different modes of regulation in different databases, these were considered twice for determining the mode of regulation to use in the CollecTRI-derived regulons. We then assigned a mode of regulation, indicating the effect (activating or repressing) of transcriptional regulation from the TF to its target gene, to each TF-target interaction based on multiple sources of information. We first checked the specific knowledge for each TF-target interaction based on the prevalence of PMIDs linked to a particular mode of regulation (activation or repression). If no information or the same amount of information about the mode of regulation was present for a TF-gene interaction, we checked information gathered about the general mode of regulation of a TF to assign an activating or repressing mode to these TF-gene interactions. We extracted regulatory information from GO terms (30) and Uniprot keywords (32), as well as the characterization and classification of effector domains of 594 human TFs provided by Soto et al. (33). More specifically we checked if the TF was annotated with either the specific term or any child term of MF_GO:0001228, MF_GO:0001217, MF_GO:0001217, MF_GO:0003713, MF_GO:0003714, BP_GO:0045944 or BP_GO:0000122. From the uniprot keywords we extracted the information on the TF role based on the UniProtKB keywords Activator (KW-0010) and Repressor (KW-0678). If all three sources, meaning the classification based on the GO terms, the Uniport keywords, and the effector domains, agreed on the regulatory effect of a TF, the mode of regulation was assigned accordingly for all target genes without any prior information on the mode of regulation from the PMIDs. In addition, we also considered information from other interactions in the regulon of a TF to classify it as activator or repressor. Hereby,, we examined all TF-gene interactions with an assigned mode of regulation in the regulon of a TF and classified a TF based on whether the majority of these interactions were linked to an activating or repressing mode of regulation. In instances where no information was available from any of the sources, we, by default, assigned the activating mode to the TF-gene interactions in question.

### RegNetwork, ChEA3, Pathway Commons and DoRothEA

RegNetwork (26) human regulons were downloaded from their website (https://regnetworkweb.org/download.jsp). The TF regulons from ARCHS4_Coexpression (ChEA3 ARCHS4), ENCODE_ChIP-seq (ChEA3 ENCODE), Enrichr_Queries (ChEA3 Enrichr), GTEx_Coexpression (ChEA3 GTEX), Literature_ChIP-seq (ChEA3 Literature) and ReMap_ChIP-seq (ChEA3 ReMap) were downloaded from their website (https://maayanlab.cloud/chea3/) (24). DoRothEA (12) was downloaded using the function get_dorothea from the decoupler v2.4.0 bioconductor package (34) and filtered by its confidence levels A, B and C. For each TF-gene interaction where the sign of regulation was not stated, an activating mode of regulation was assigned by default.

### Computing TF-gene weights

We employed two different tools, namely *MatrixRider* (35) and *FIMO* (36), to perform motif enrichment analysis and calculate binding weights for the TF-gene interactions in the CollecTRI-derived regulons. Specifically, we used the *Matrixrider v1*.*30*.*0* and *memes v1*.*6*.*0* bioconductor packages. Before running the methods, we extracted 1,000 base pairs (bp) upstream and 100 bp downstream of the transcription start site of each gene (TSS), defining the promoter region, as well as 10,000 base pairs (bp) upstream and 100 bp downstream of the TSS, reflecting proximal regulatory regions using the

*TSS*.*TxDb*.*Hsapiens*.*UCSC*.*hg38*.*knownGene v3*.*4*.*6* package (37). Human TF binding motifs were downloaded from *MotifDb v1*.*40*.*0* (38). TF-gene pairs for which either the promoter sequence or the TF binding motif were not available were removed from the network. For the remaining 40,440 TF-gene pairs the two different tools were used as follows. MatrixRider was used to calculate binding weights for each TF-gene interaction as described in their reference manual. TF binding motifs were provided as position frequency matrices, DNA sequences of the target genes as DNAString objects and a cutoff parameter of 0 were passed to the getSeqOccupancy function within the *Matrixrider v1*.*30*.*0* package. For FIMO, TF binding motifs and DNA sequences were passed to the runFimo function within the *memes v1*.*6*.*0* package and the highest score was kept as the binding weight. For both methods, calculated binding weights were shifted to positive values with a pseudo count of 1 and normalized. We used two different normalization strategies. First, we normalized the weights per TF, meaning that the weights for all targets of a specific TF were divided by the highest TF-gene binding weight of that TF. Secondly, we normalized the weights per gene, meaning that the weights for all TFs regulating a specific gene were divided by the highest TF-gene binding weight of that gene. The final weights were compared with each other using Pearson correlation. The benchmark procedure was then repeated for the weighted network, using the calculated weights from MatrixRider with a window frame of 1,000 bp before normalization, and compared to the non-weighted CollecTRI regulons. Additionally, TF-gene links with binding weights in the lowest 10, 20 and 30% quantile were removed from the network and their performance evaluated in the benchmark.

### Benchmark data

Differentially expressed gene tables and meta data were downloaded from 907 manually curated RNA-seq and microarray experiments, collected in knockTF (28) (http://www.licpathway.net/KnockTF/download.php). These datasets include knockdown/knockout experiments across multiple tissues and cell types associated with 456 different disrupted TFs. Perturbation experiments with a perturbed TF’s log fold change greater than -1 were excluded from the final benchmark set, leading to 388 data sets covering 234 unique perturbed TFs. For each resource only perturbation experiments of TFs covered in that network were used for the benchmark (Table 1).

**Table 1.**
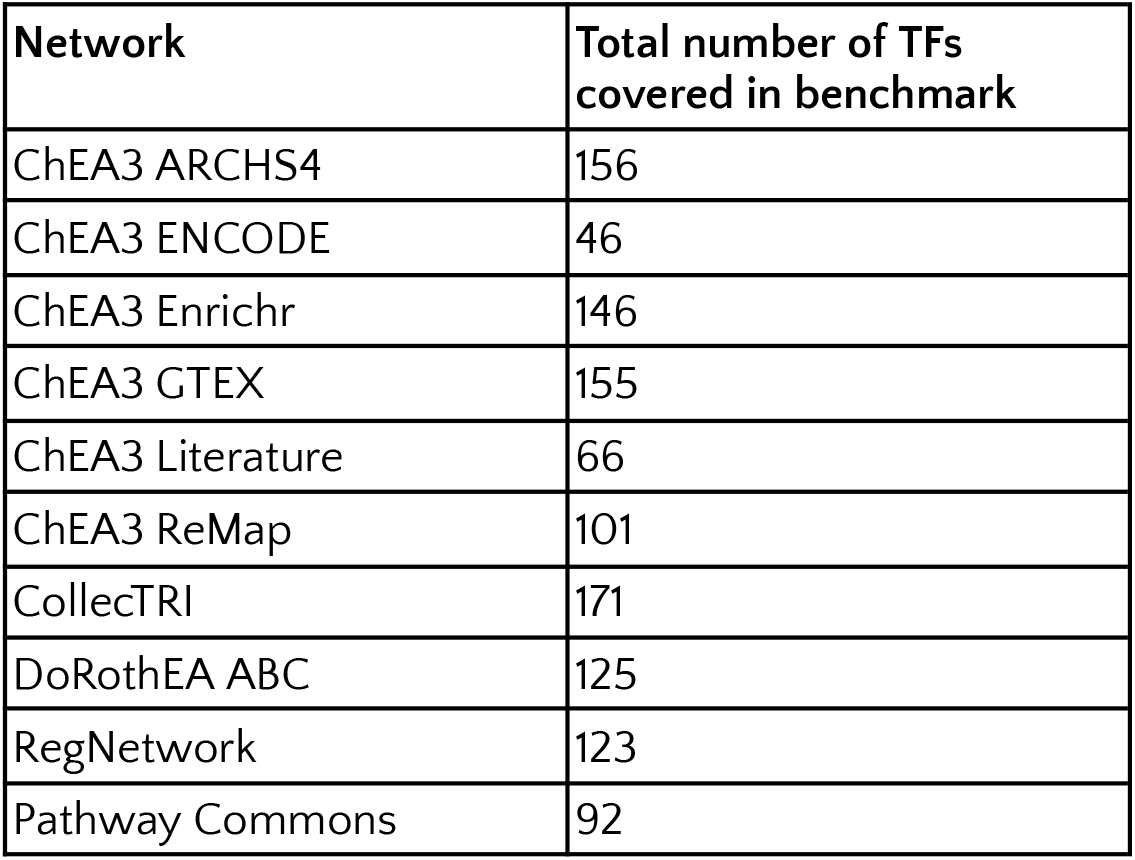
Overview tested TFs per network.

| Network | Total number of TFs covered in benchmark |
| --- | --- |
| ChEA3 ARCHS4 | 156 |
| ChEA3 ENCODE | 46 |
| ChEA3 Enrichr | 146 |
| ChEA3 GTEx | 155 |
| ChEA3 Literature | 66 |
| ChEA3 ReMap | 101 |
| CollecTRI | 171 |
| DoRothEA ABC | 125 |
| RegNetwork | 123 |
| Pathway Commons | 92 |

### TF activity estimation

TF activities were estimated based on the log fold-changes of the direct target genes after perturbation. Each network was first filtered to keep only TF-gene interactions of genes measured in the experiment. We then selected TFs with at least five gene targets and estimated TF activities using the consensus score of the multivariate linear model method, the univariate linear model method and the weighted sum method using the *decoupler v1*.*2*.*0* python package (34).

### Benchmark procedure

The benchmark was performed using the benchmark function from the *decoupler v1*.*2*.*0* python package (34). To globally evaluate collections of TF regulons, TF activity scores obtained as described above were first multiplied by the sign of the perturbation (knockout: negative, overexpression: positive) for each perturbation experiment. The activity scores matrix (rows: experiments, columns: TF activities) is then flattened across experiments into a single vector. The objective is to distinguish between perturbed TFs, the true positives, from all unperturbed ones, true negatives. However, due to the large class imbalance between true positives and negatives, a downsampling strategy was employed within the benchmark. For each permutation, an equal number of positive and negative classes were randomly selected 1,000 times to calculate the area under the Receiver Operating Characteristic (AUROC) and Precision-Recall Curve (AUPRC) metrics, obtaining distributions of prediction measurements.

The performance evaluation for specific TFs was performed only for TFs for which at least 5 experiments were available in the KnockTF dataset (after filtering out experiments with insufficient perturbation, Methods 4.6). In this setting, the objective is to distinguish between perturbed experiments for each retained TF, the true positives, from all the unperturbed ones, true negatives. The same strategy as described above is used, but instead of flattening the activity scores matrix, only the vector of one TF is extracted for evaluating the performance. This was done separately for each of the TFs in Supplementary Figure 2.

### Evaluation of size bias

For the three top performing regulon collections, namely CollecTRI, DoRothEA ABC and RegNetwork, we used a two-sided t-test which was adjusted for multiple testing using Benjamini-Hochberg correction to compare if there was a difference in the number of targets for TFs that were part of the benchmark data set, compared to the TFs that were not. Pearson correlation coefficients were computed to assess the relationship between the number of targets and the absolute activity scores of TFs across all benchmark experiments included in the benchmark. We then summarized the correlation between the absolute scores and the number of targets across experiments with the mean correlation and compared it across networks.

### Case study

We tested the network by estimating TF activities in three types of cancer: Uterine Corpus Endometrial Carcinoma (UCEC), Lung Adenocarcinoma (LUAD), and Clear Cell Renal Cell Carcinoma (CCRCC), retrieved from the third phase of the Clinical Proteomic Tumor Analysis Consortium (CPTAC). The data was provided by Gaytan et al. (in preparation) which was collected from NCI’s Genomic Data Commons (GDC) (39) using the GDC transfer tool. Raw count tables were subjected to VSN normalization, after filtering genes with a low number of counts (40). For each cancer type, we then performed differential expression analysis between tumor and natural tissue samples using the limma R package (41). The t-values were used to estimate the consensus TF activity as described previously. A significance threshold of 0.05 was applied.

## RESULTS

### Sources of prior knowledge on TF-gene regulatory interactions

CollecTRI (Collection of Transcription Regulation Interactions) is a compilation of available transcription regulation information from databases integrated with information extracted from the text-mined resource ExTRI (23), which extracts sentences containing descriptions of transcription regulation events, known as TRIs (Transcription Regulation Interactions). Here, we used those resources after updating some of the databases and including three new ones (Table 2): Pavlidis (42), NTNU Curated (43) and DoRothEA A (12) (Supp. Figure 1). NTNU Curated are a subset of ExTRI sentences manually curated for validity of TF-gene interaction and mode of regulation, and DoRothEA A contains the TF-gene interactions with the highest confidence level of the DoRothEA meta-resource (12), which is also evaluated in this publication separately. Note that DoRothEA compiles some of the same resources as CollecTRI; however, the overlap is taken into account for the creation of the CollecTRI-derived regulons as it is based on the number of unique PubMed IDs (PMIDs) supporting each annotation across resources.

**Table 2.**
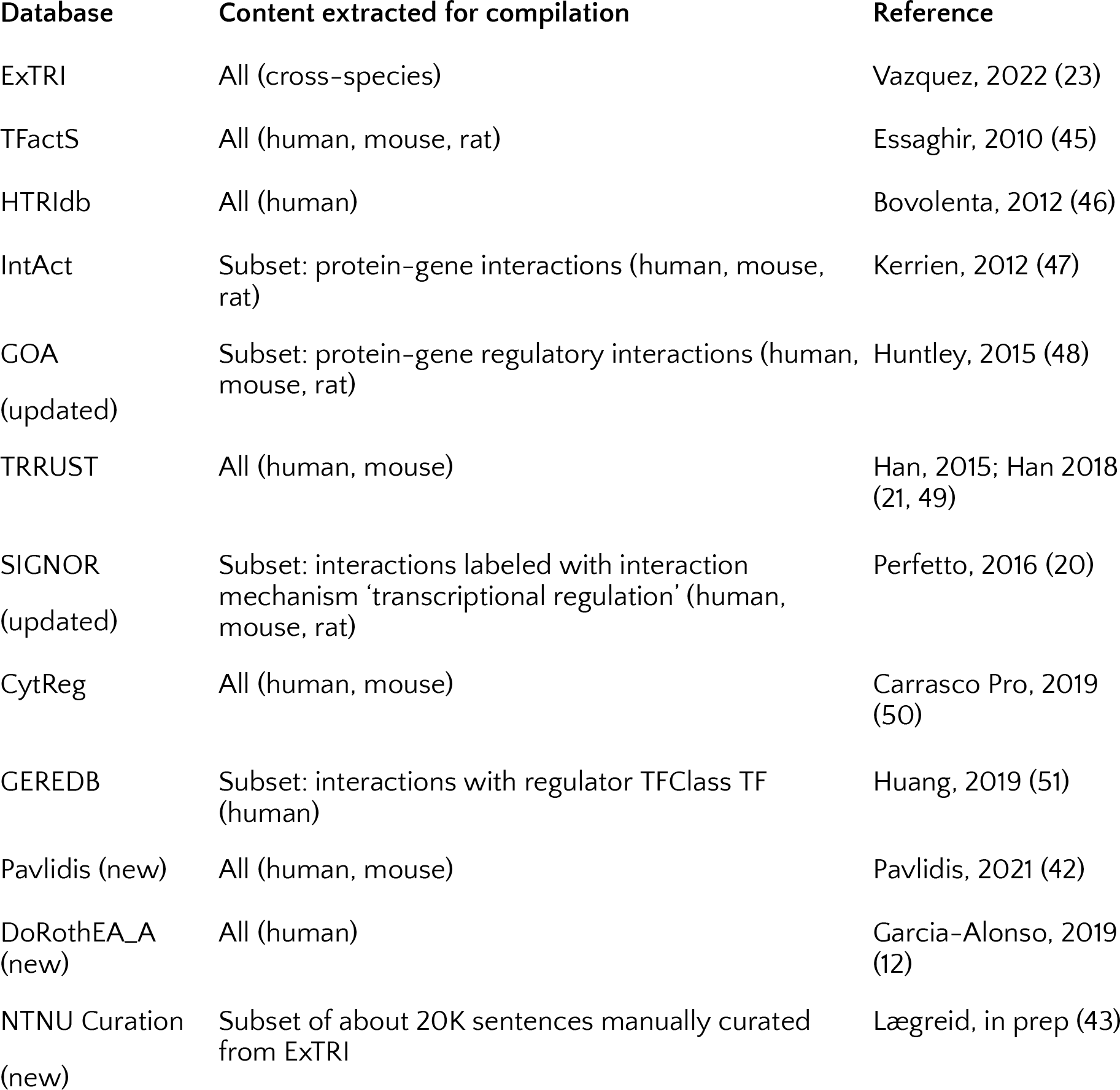
Resources of TF-gene interactions used in CollecTRI. The table indicates whether all interactions or subsets of them were included in CollecTRI.

| Database | Content extracted for compilation | Reference |
| --- | --- | --- |
| ExTRI | All (cross-species) | Vazquez, 2022 (23) |
| TFactS | All (human, mouse, rat) | Essaghir, 2010 (45) |
| HTRIdb | All (human) | Bovolenta, 2012 (46) |
| IntAct | Subset: protein-gene interactions (human, mouse, rat) | Kerrien, 2012 (47) |
| GOA<br>(updated) | Subset: protein-gene regulatory interactions (human, mouse, rat) | Huntley, 2015 (48) |
| TRRUST | All (human, mouse) | Han, 2015; Han 2018 (21, 49) |
| SIGNOR<br>(updated) | Subset: interactions labeled with interaction mechanism 'transcriptional regulation' (human, mouse, rat) | Perfetto, 2016 (20) |
| CytReg | All (human, mouse) | Carrasco Pro, 2019 (50) |
| GEREDB | Subset: interactions with regulator TFClass TF (human) | Huang, 2019 (51) |
| Pavlidis (new) | All (human, mouse) | Pavlidis, 2021 (42) |
| DoRothEA_A<br>(new) | All (human) | Garcia-Alonso, 2019 (12) |
| NTNU Curation<br>(new) | Subset of about 20K sentences manually curated from ExTRI | Lægreid, in prep (43) |

From the resources, all instances of genes or proteins have been translated into human gene symbols, including mentions to rat or mouse entities which were translated with the help of orthology tables. We decided to also consider TF-gene interactions from mouse and rat, as it is a common practice in the field to assume that the TRIs translate across murine organisms and humans due to the high conservation of regulatory mechanisms across these organisms (44). Additionally, a large component of CollecTRI is extracted from text-mining, where it is often difficult to assign the correct species to information extracted from PubMed abstracts; in fact, this information may be missing entirely from abstracts.

Moreover, two TF dimers, AP1 and NFKB, were treated as transcription factors themselves in CollecTRI since they, in the literature, are very frequently mentioned only by their dimer name. When merging the CollecTRI resources, the information regarding the monomer AP1 or NFKB constituents (e.g. JUN- or FOS-family or NFKB1) was merged into information referring to the complex (i.e. AP1 for JUN and FOS) and vice versa, see details in (23).

### Construction of TF regulons from CollecTRI

From the compiled information in CollecTRI we constructed signed and directed CollecTRI-derived regulons which can be used for the inference of TF activities. To ensure reliability of the TF-gene interactions and account for the overlap across resources, we gathered the unique PMIDs for each TF-gene interaction and removed those lacking any reference (Figure 1A). Furthermore, to focus on proteins with a direct regulatory effect on gene expression, we included only TF-gene links from TFs classified as DNA-binding transcription factors (dbTFs), co-regulatory transcription factors (coTFs) or general initiation transcription factors (GTFs) based on criteria from TFclass (52), Lambert et al. (53), Lovering et al. (6), or gene ontology (GO) annotations (30).

We then assigned a mode of regulation to each TF-gene pair, indicating the sign of transcriptional regulation from the TF to its target gene. Specifically, we determined whether the TF activates or represses the expression of its target gene (Figure 1B). Hereby, activation corresponds to an increase in the expression of the target gene, whereas repression corresponds to a decrease in expression. To determine the sign of transcriptional regulation for each TF-gene pair in the CollecTRI-derived regulons, we integrated information from multiple resources, including the specific knowledge for each TF-target link based on the prevalence of PubMed references linked to a particular mode of regulation. We also incorporated general prior knowledge about the mode of regulation of a TF derived from GO terms (30) and Uniprot keywords (32), as well as the classification of effector domains of 594 human TFs provided by Soto et al. (33). Given that the effector domain can activate or repress the expression of a TF’s target genes through several mechanisms, it provides additional information for the classification of TFs into activators or repressors (54). Furthermore, we considered information from other interactions in the regulon of a TF to make a decision on whether a TF is more likely to activate or repress the expression of its target gene. The information from each source was considered separately, and the prevalence of PMIDs was prioritized to determine the mode of regulation. If information from PMIDs was not available, we relied on the general prior knowledge about the mode of regulation of a TF and lastly on the information about other interactions in the TFs regulon to select the mode of regulation for all TF-gene pairs in the GRN generated based on CollecTRI (Methods: Assigning the TF-gene mode of regulation). In instances where no information was available from any of the sources, we designated TFs as activators by default, and assigned a mode of regulation accordingly (Figure 1B). For 18,345 TF-gene interactions, the mode of regulation was determined based on the prevalence of PubMed references. Meanwhile, for 9,226 and 949 interactions, the mode of regulation was assigned based on the prior knowledge of the TFs and other interactions in the regulon, respectively. For the remaining 17,364 TF-gene interaction an activating mode was assigned by default (Supp. Figure 2A). This annotation procedure led to 83% activatingand 17% repressing TF-gene links. 51% of TFs were represented with a dual role in regulation, meaning the TF was assigned to either activate or repress the expression of its target genes. 33% of the TFs had only activating links, whereas 16% of TFs are only represented by repressing links. (Figure 1C). In total, the CollecTRI-derived regulons cover 1,183 TFs and 45,856 signed TF-gene interactions.

**Figure 1.**
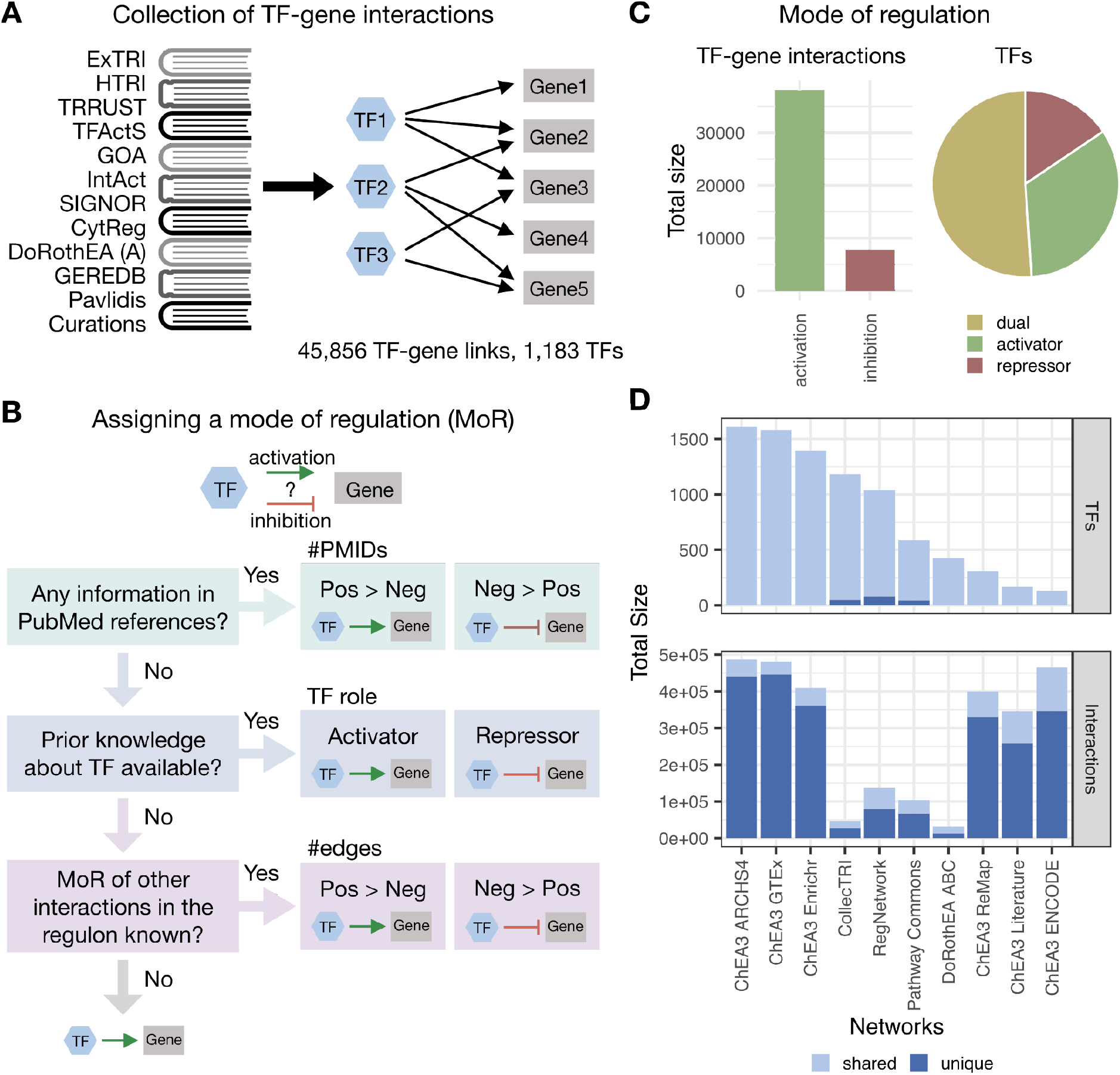
Description of transcription factor (TF)-gene interactions in the CollecTRI-derived regulons and comparison to other regulon collections. (A) Collecting transcription factor (TF)-gene links to construct regulons from CollecTRI. Depicting prior knowledge resources used to collect links, which were aggregated within CollecTRI. (B) Flow chart describing how the mode of regulation (MoR) was assigned to each TF-gene link. The MoR, indicating the direction of transcriptional regulation from the TF to its target gene, was determined for each TF-gene link, based on factors such as PubMed references (PMIDs), prior classification of the TF and the MoR of other genes in the regulon. (C) Summary of the MoR for TF-gene interactions in CollecTRI. Total number of interactions for activating and repressive TF-gene links (left) and percentage of TFs that purely function as activators, repressors, or have a dual mode of regulation (right). (D) Comparison of the number of unique TFs (top) and interactions (bottom) across different resources - with ChEA3 ARCHS4, ChEA3 GTEx and ChEA3 Enrichr being solely based on co-expression or co-occurrence. Any TF or interaction present in more than one resource is considered shared.

We compared the coverage of the CollecTRI-derived regulons to other known collections of TF regulons, namely ChEA3 (24), RegNetwork (26), Pathway Commons (25) and DoRothEA (12). ChEA3 contains a collection of gene set libraries generated from TF-gene co-expression, TF-target associations from ChIP-seq experiments, and TF-gene co-occurrence computed from user-submitted lists to the Enrichr tool. RegNetwork is a manually curated database of experimentally observed or predicted transcriptional and post-transcriptional regulatory interactions. Pathway Commons is a resource that compiles information about regulatory networks as well as biological pathways including molecular interactions, signaling pathways, and DNA binding from different databases. Finally, DoRothEA integrates information on gene regulatory interactions with assigned confidence levels from multiple sources, including literature-curated resources, ChIP-seq peaks, motif analysis, as well as inference from gene expression data. Only DoRothEA among the four regulon collections we compared to the CollecTRI-derived regulons also provides signed information about the direction of transcriptional regulation. For a fair comparison between the TF regulons, all collections were filtered to only contain TF-gene interactions from annotated dbTFs, coTFs, or GTFs, as described above.

We then compared the TFs and TF-gene links across all collections of TF regulons, and we found that the CollecTRI-derived regulons exhibit the highest TF coverage (1,183) besides the ChEA3 gene set libraries ARCHS4 (1,612), GTEx (1,578) and Enrichr (1,393). It is worth noting that these ChEA3 libraries were generated using co-expression or co-occurrence strategies, which are known to produce a higher number of false positive interactions in TF-target association studies (27). The CollecTRI regulons cover 47 TFs not present in any of the other four resources. RegNetwork and Pathway Commons also provide information on 80 and 42 unique TFs, respectively, otherwise 91.3% of all TFs across the analyzed resources are present in at least two of them. In terms of TF-gene interactions, resources mainly collecting information from curated databases, such as RegNetwork, Pathway Commons, DoRothEA and CollecTRI, generally showed a lower number of interactions. As previously mentioned, TF regulons generated using co-expression and co-occurrence strategies, such as some ChEA3 libraries, tend to have a higher number of potential interactions that often include many indirect regulatory relationships. In general, there was a low overlap between the resources we compared, with an average of 72.4% of interactions being unique to each collection of TF regulons (Figure 1D). Overall, the CollecTRI-derived regulons have an extensive coverage of TFs with high-confidence interactions and, in contrast to most other regulon collections, include information about the sign of the transcriptional regulation.

### Systematic comparison of TF activity inference from CollecTRI-derived regulons with other regulon collections

We evaluated the quality of the CollecTRI-derived regulons by assessing how well they are able to recapitulate the changes in gene expression caused by the perturbation of a TF in comparison to other existing regulon collections. As previously described, we reasoned that if a TF’s set of targets is reliable, meaning their expression is directly regulated by the TF, the regulon’s collective expression pattern should be a proxy of the TF’s transcriptional activity (12). To test this, we downloaded perturbation data from KnockTF (28), a comprehensive human gene expression profile database from TF knockdowns and knockouts studies. KnockTF contains manually curated RNA-seq and microarray datasets associated with TFs perturbed by different knockdown or knockout techniques across multiple tissues and cell types. For the benchmark, we restrict the datasets to experiments were the TF perturbation is highly likely to have been effective, by only including data from experiments where the expression of a TF was markedly decreased after its knock down or knock out, leading to a total number of 388 perturbation experiments covering 234 unique TFs (Methods: Benchmark data).

We then followed the benchmark pipeline in the decoupler python package (34) to systematically compare the regulons generated from CollecTRI, to the ones from DoRothEA, Pathway Commons, RegNetwork and the ChEA3 libraries. Additionally, we used a permuted version of the CollecTRI-regulons as a baseline of performance. In this version the target genes and mode of regulation in CollecTRI were shuffled and randomly assigned to a TF. As such, these TF regulons do not represent biological information and can thus serve as a baseline of performance. TF activities were then inferred from the differentially expressed genes of each KnockTF experiment using the regulons provided by each resource. Only TF regulons containing at least five target genes among the genes measured in the experiment were used for the activity inference, leading to a restricted number of TFs for each resource (Supp. Figure 3). All inferred TF activities across experiments were sorted by their activity scores, and the classification of TFs based on the estimated activities compared to the knock-out information was evaluated with the area under the receiver operating characteristic curve (AUROC) and the area under the precision-recall curve (AUPRC) (Figure 2A) (Methods: Benchmark procedure). Before systematically comparing CollecTRI regulons to those from other resources, we first evaluated the effect of assigning the sign of regulation to the TF-gene interactions. We found that the fully signed CollecTRI-derived TF regulons perform better in the benchmark than TF regulons with the mode of regulation for all TF-gene rendered activating (adjusted p-value < 2.2×10^−16^, mean t-value across tests equal to 64.9 and 51.6 for AUROC and AUPRC, respectively) (Supp. Figure 2B). We then performed the comparison of CollecTRI-derived regulons to those from other resources and showed that the CollecTRI regulons had median AUROC and AUPRC values of 0.7 and 0.75, respectively, which were higher than those of all other resources (adjusted p-value < 2.2×10^−16^, mean t-value across tests equal to 252 and 275.1 for AUROC and AUPRC, respectively) (Supp. File 1, Figure 2B). Furthermore, all ChEA3 libraries, except for ChEA3 ARCHS4, did not exhibit a higher performance compared to the random baseline set by the permuted CollecTRI version (t-test: adjusted p-value > 0.05). Overall, the results from the benchmark show that the CollecTRI-derived regulons outperforms other TF regulon collections in identifying perturbed TFs based on TF activities, suggesting that, of the resources here compared, the TF-gene interaction information compiled in CollecTRI provides the most reliable starting point for estimating TF activities.

**Figure 2.**
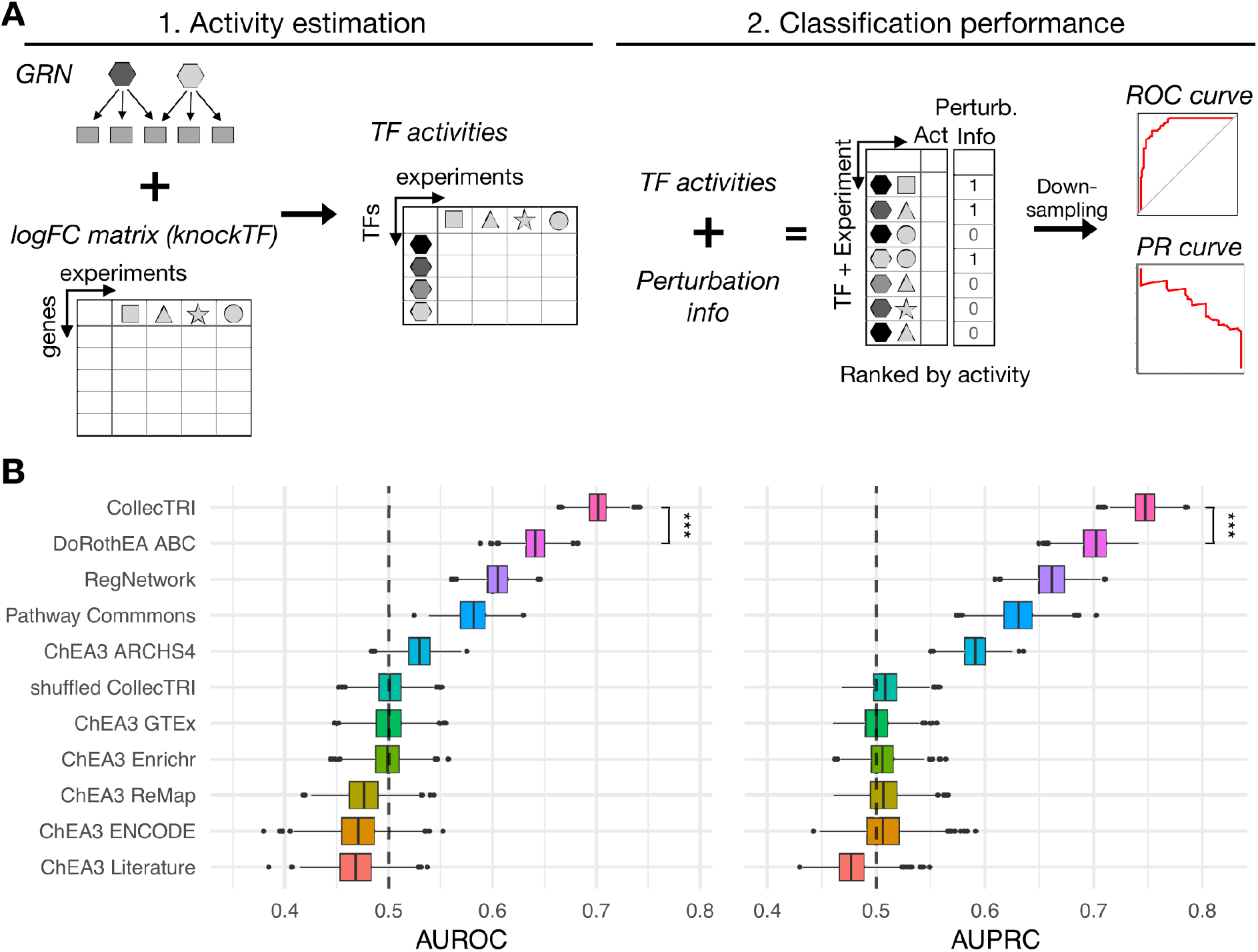
Systematic comparison of collections of transcription factor (TF) regulons. (A) Description of benchmark pipeline for the comparison of different regulon collections. First, transcription factor (TF) activities are inferred from the gene expression data of the knockTF perturbation experiments using the regulon information from each resource. TFs are presented as differently colored hexagons and experiments are presented as different shapes. Activities are then aggregated across experiments and ranked by their activity. A downsampling strategy is applied to have an equal number of perturbed and non-perturbed TFs randomly selected 1,000 times to calculate area under the Receiver operating characteristic (AUROC) and Precision-Recall curve metrics (AUPRC). (B) Predictive performance of TF regulons identifying perturbed TFs in knockTF experiments. AUROC (left) and AUPRC (right) for each regulon collection classifying TFs as perturbed or non-perturbed based on their activities.

Since the benchmark data mainly covers TFs that are well studied and usually have a larger number of targets associated with them, we tested if the number of genes regulated by a TF was related to the performance of the networks to predict perturbed TFs. For the top three performing TF regulon collections, we first tested if there was a difference in the number of targets between TFs that were part of the benchmark data set and those TFs that were not. For all three resources, we observed that the TFs included in the benchmark had a higher number of targets associated with them (adjusted p-value = 1.76^-5^, 1.34^-3^ and 2.8^-4^, t-value = 4.68, 3.27 and 3.84 for CollecTRI, DoRothEA and RegNetwork, respectively) (Supp. Figure 4A). To assess the relationship between the number of targets and the accuracy estimating TF activities for each experiment included in the benchmark, we computed Pearson correlation coefficients and found that the average correlation across all experiments was equal or less than 0.3 for all resources, with the CollecTRI-derived regulons showing the lowest mean correlation of 0.14 (Supp. Figure 4B). Therefore, we concluded that the better performance of the CollecTRI-derived regulons is not influenced by an increased bias towards TFs with a higher number of targets.

Another limitation of the current benchmark is that it disregards possible off-target effects of TF perturbation assuming that the perturbed TF has the most deregulated activity. Thus for a limited collection of 12 TFs where we had multiple perturbation experiments we repeated the benchmark only classifying the activity of the perturbed TFs without including non-perturbed TFs (Methods: Benchmark procedure). In this benchmark setting, we observed a better performance of CollecTRI regulons for the TFs REST, TP53, FLI1, NRF2F2 and SOX2 in comparison to the other networks with average median AUROC and AUPRC value of 0.84 and 0.88, respectively, (adjusted p-value < 0.001, mean t-value across TFs = 77.6 and 76.4 for AUROC and AUPRC, respectively) and a perfect classification for REST (Supp. Figure 5A). However, overall all networks performed comparably (Supp. Figure 5B). Although limited to a few TFs, CollecTRI-derived regulon’s performance was comparable to the other networks in this benchmark setting, with an improved performance for specific TFs.

### Case study

To showcase the value of using CollecTRI-derived GRN for predicting TF activities, we performed a TF activity inference analysis using differential expression data from three cancer types: Uterine Corpus Endometrial Carcinoma (UCEC) (55), Lung Adenocarcinoma (LUAD) (56), and Clear Cell Renal Cell Carcinoma (CCRCC) (57) (Figure 3A). These datasets comprise gene expression data of tumors and adjacent normal tissues from multiple patients.

**Figure 3.**
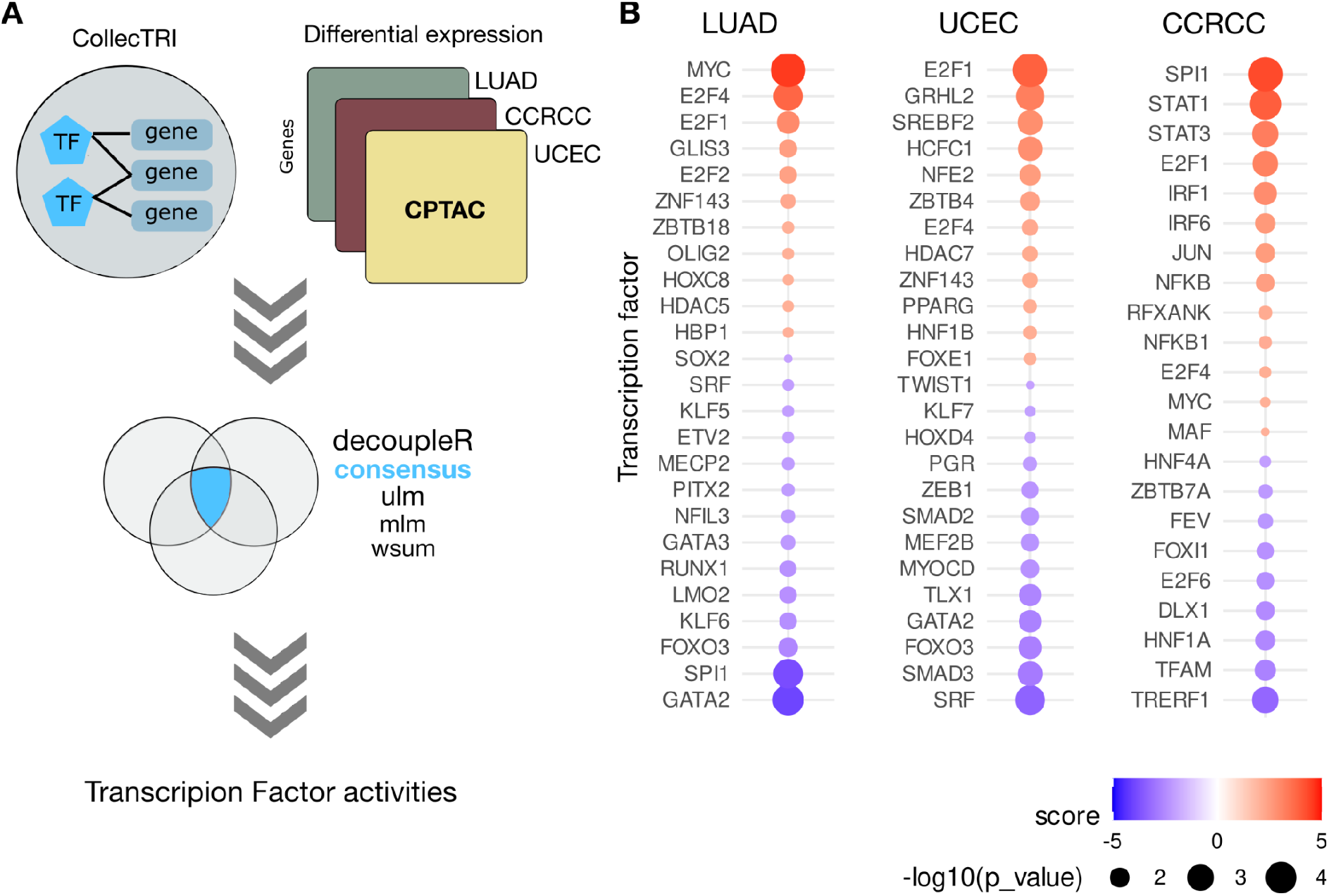
Workflow and results of the case study of transcription factor activity inference using CollecTRI and decoupleR. (A) Schematic representation of the workflow for the inference of transcription factor activities. (B) Transcription factors with differential activities as predicted by decoupleR. Each TF is shown as an individual dot and colored based on the consensus score. The size of the dot is inversely related to the p-value (the bigger the size, the more significant the observation). Abbreviations: TF: Transcription factor, Uterine Corpus Endometrial Carcinoma (UCEC), Lung Adenocarcinoma (LUAD), Clear Cell Renal Cell Carcinoma (CCRCC).

Based on the differentially expressed transcriptome of tumor versus normal tissue, we predicted TF activities for each cancer type and observed in total 62 significantly deregulated TFs, as shown in Figure 3B. Our analysis generally reflected previously described TF activity changes in cancer tissue. For instance, proliferation- and cell survival-promoting TFs, such as MYC, Jun-, FOS-, and E2F family TFs, were found to have significantly increased activity across the three cancer types. On the other hand, cell death-related TFs, such as members of the FOXO family, were found to have significantly reduced activity (Supplementary Table 2).

To highlight the added value of the additional TF-gene coverage of CollecTRI, we compared the predictions of TF activities for those TFs whose regulons are included only in CollecTRI-derived GRN and not in DoRothEA ABC, which was chosen as the main network for comparison as it was the second-best performing network in our benchmark. To evaluate the validity and relevance of the “CollecTRI-exclusive” TFs, a literature review for their role in their respective cancer types was conducted.

For LUAD, in total six TFs were uniquely part of CollecTRI regulons, and four of those had a previously reported role in several aspects of LUAD, such as its development and prognosis. OLIG2 and ETV2 are TFs reported to be overexpressed in lung adenocarcinomas (58, 59). Similarly, the upregulation of HDAC5 has been found to promote lung adenocarcinoma by regulating several cell cycle and epithelial-mesenchymal transition genes (60), and in our analysis, CollecTRI regulons predicted its increased activity in LUAD. LMO2 is a tumor suppressor which acts through the regulation of the Wnt pathway in several tumor types. In lung adenocarcinomas and other epithelial-derived tumors, LMO2 was found to have a reduced expression and activity (61), as also predicted in our results.

Among the TFs for which we estimated altered activities in UCEC, eight were part of only CollecTRI-regulons, four of which had a previously described role in this specific cancer type. SMAD2, together with SMAD3, has been shown to have tumor-suppressive functions in endometrial carcinoma cells, and the inhibition of its activity has been associated with the constituent activation of the PI3K/AKT pathway, increased proliferation and decreased apoptosis (62). Both SMAD3 and SMAD2 showed a significantly decreased activity in the UCEC dataset, with the latter being part uniquely of CollecTRI regulons. As the afore-described HDAC5, HDAC7 is another histone deacetylase, whose inhibition with pan-HDAC inhibitors has been reported to lead to cell cycle arrest in UCEC (63), suggesting a positive role of the TF in cancer development, which is also reflected in the increased activity predicted with CollecTRI regulons. The HCFC1 transcription factor has an immunomodulatory role in cancer by inhibiting immune responses, and by promoting tumor growth and vascularization (64). In accordance with its cancer-promoting role, HCFC1 was found to have an increased activity in UCEC. The ZBTB4 TF, a proposed tumor suppressor, was the only instance where CollecTRI activities did not correspond to the expected activity given the role of the TF. ZBTB4 is an essential component in maintaining genomic stability (65), and its expression is decreased in endometrial cancer (66). At the same time, higher expression of ZBTB4 has been proposed as a favorable marker of relapse-free survival in other types of cancers, such as breast cancer (67). ZBTB4 was found to be overactivated in the studied UCEC samples, suggesting that additional validation is needed to evaluate the prediction.

CCRCC is one of the three main subtypes of renal cell carcinomas (RCCs), all of which have been described for their distinct transcriptional and epigenetic characteristics. A study by (68) reported the main driving TF of each subtype. Among those main driving TFs, two are found in the predictions by CollecTRI-regulons. ETS1, which was estimated to be overactive by our analysis, was found to be one of the two main TF regulators in CCRCC. On the contrary, FOXI1, a main TF regulator of another RCC, chromophobe RCC, was found to be significantly underactive in CCRCC, as would be expected in this specific subtype. FOXI1 was among the TFs which were exclusively found with ColleCTRI regulons. TFAM is a mitochondrial TF which is also included in the regulation of pyroptosis. Together with 10 more pyroptosis-related genes, TFAM, was identified as a risk gene for the prognosis of CCRCC (69). KLF7’s exact role in CCRCC is not clear, however, KLF7 serves as a target for miR-22 which has been suggested as an important regulator and prognostic marker for CCRCC (70). Furthermore, according to The Human Protein Atlas, its high expression has been correlated with a favorable prognosis for kidney carcinomas. Two additional TFs, TRERF1 and DLX1 were inferred to be more active in tumor than normal tissue. While the role of TRERF1 in CCRCC is poorly understood, TRERF1 is a known regulator of CYP11A1, which is frequently downregulated in CCRCC. DLX1 has no reported role in CCRCC, however, it has a known oncogenic role in other cancer types such as prostate (71) and ovarian (72) cancers.

Overall, this comparative analysis highlights the usefulness of CollecTRI-derived regulons in inferring TF activities. The presence of cancer-type-relevant TFs in the results showcases how the augmented TF coverage in CollecTRI-derived regulons appears to balance the identification of meaningful TFs without overwhelming the output with potentially extraneous information.

## DISCUSSION

Transcription factor (TF) regulons represent regulatory circuits that depict the coordinated regulation of downstream target genes by TFs. They can be valuable for understanding various biological processes, including development, cell differentiation, tissue homeostasis, and disease progression. To derive functional insights from these regulons, TF activities can be inferred from the expression levels of target genes, as shown in various studies (9, 73–75). However, to interpret these findings accurately, it is important to critically evaluate the reliability and coverage of TF regulons.

In this paper, we present a well-defined, transparent, and reproducible workflow to generate regulons from CollecTRI, a meta-resource that compiles TF-gene information from 12 different resources including information inferred from text mining, manual curations and a number of publicly available databases (23). With that, the CollecTRI-derived regulons provide the most extensive coverage of TF-gene interactions compared to other collections of regulons that extract TF-gene interaction knowledge solely from literature. Since most publicly available meta-resources of TF-gene interactions contain limited or no information about the mode of regulation of a TF to its target genes, we propose an evidence-driven approach to infer the sign of regulation for each TF-gene link in the CollecTRI regulons, which can also be applied to other comprehensive knowledge bases. We evaluated the approach and confirmed that adding the information about the sign of regulation to the CollecTRI regulons leads to more accurate TF activity inference. Next, through systematic comparison with other known TF regulon collections, we showed that CollecTRI-derived regulons perform best in identifying perturbed TFs based on gene expression, suggesting a high quality in CollecTRI’s TF-gene interactions. Finally, we showcase the value of the CollecTRI regulons in inferring TF activities in three different cancer types and successfully identifying changes in the activity levels of TFs known to be involved in these contexts.

Despite the good performance of the CollecTRI regulons in the systematic comparison, it is important to bear in mind that the current benchmark is limited to a specific set of TFs. Further perturbation studies would therefore be useful to extend the current benchmark and allow for a more comprehensive evaluation of CollecTRI and other resources.

While the coverage in the CollecTRI regulons is substantially larger than those of other resources, it could still be expanded by including additional TF-gene interactions from other resources. However, identifying high-quality TF-gene interactions within a resource and distinguishing them from indirect regulatory relationships is challenging. Since CollecTRI is primarily assembled from literature-curated resources, a bias for well-studied TFs may be present. We observed similar bias trends across meta-resources, quantified as the correlation of inferred TF activities with the number of targets of each TF.

Another limitation is that the CollecTRI regulons currently only take the sign of regulation into account, omitting the quantitative nature of gene regulation (4). We therefore estimated TF binding weights using motif enrichment analysis, but observed no benefit in the inference of TF activities (Supplementary Note 1, Supp. Figure 6). Since CollecTRI compiles exclusively TF-gene link interactions omitting cooperative events between TFs and other proteins, distal interactions and the chromatin accessibility landscape among other processes (4), it only captures one layer of the cis-regulatory code. This might explain why using TF binding weights did not increase the overall predictability of perturbed TFs.

Finally, the CollecTRI regulons were constructed as generalistic interactions and, as such, do not account for cell type-specific differences (76). Nonetheless, CollecTRI regulons could be used as a building block for context-specific interactions using complementary data types, such as single-cell transcriptomic or chromatin accessibility data.

In summary, we constructed a collection of TF regulons with a high coverage of TFs and high confidence TF-gene interactions, which is freely available to the community via OmniPath (29) and DoRothEA (12) packages. We conducted a systematic comparison with other known resources, where the CollecTRI regulons showed the best performance in recapitulating changes in gene expression caused by the perturbation of a TF. Additionally, we demonstrated how the regulons can be applied in a biological context and can help uncover the role of transcriptional regulation in various biological contexts.

## DATA AVAILABILITY

The code for the curation of regulatory interactions of CollecTRI and the construction of the CollecTRI-derived regulons is available here: https://github.com/Rbbt-workflows/ExTRI, https://github.com/saezlab/CollecTRI. Files necessary to reproduce the presented results are can be downloaded from https://zenodo.org/record/7773985#.ZCSMiexBw-Q. The CollecTRI-derived regulons are freely available and can be accessed through OmniPath (https://omnipathdb.org/) (29) and the DoRothEA package (https://saezlab.github.io/dorothea/) (12), which are also available in Bioconductor (https://bioconductor.org/packages/release/bioc/html/OmnipathR.html, https://bioconductor.org/packages/release/data/experiment/html/dorothea.html).

## AUTHOR CONTRIBUTIONS

SMD constructed the CollecTRI-derived regulons and performed the evaluation and comparison. ET performed the case study. MV built the CollecTRI meta-resource. RORF advised the analysis, with input from PBiM. PBiM and RF contributed the benchmark implementation; RF reviewed the code. AL and JSR supervised the project. SMD, ET and MV wrote the manuscript, which has been revised by all authors.

## ACKNOWLEDGEMENTS

We thank Estefani Gaytan for sharing the processed CPTAC data with us prior to submission and Denes Turei for his help with the implementation of the CollecTRI-derived regulons into Omnipath.

