## Supplementary Data for "Expanding the coverage of regulons from high-confidence prior knowledge for accurate estimation of transcription factor activities"

Supplementary Note 1 ***Weighting CollecTRI interactions based on binding probabilities***

To enrich the biological information incorporated within CollecTRI, we implemented a weighting scheme for TF-gene interactions, reflecting the likelihood of a TF binding to the gene's genomic location. The underlying assumption is that a TF's activity is primarily revealed by the coordinated expression of genes where the TF has a strong binding affinity to the promoter region or other regulatory elements. Thus, genes whose genomic location is enriched by the binding motif of a TF of interest should have a higher weight in the inference of its activity, than genes in the regulon without an enriched binding motif. For that, we estimated binding weights for TF-gene pairs with known binding motifs and genomic regions using FIMO (1) and MatrixRider (2). We first inferred binding weights to the promoter regulatory region of each gene, including 1,000 base pairs (bp) upstream and 100 bp downstream of the transcription start site (TSS) of the gene, as previously proposed by Sanghi et al. (3). To also include proximal regulatory regions of the gene, we next extended the window frame to 10,000 bp upstream of the TSS. We then calculated the Pearson correlation of the inferred binding weights across the tools, window sizes, and different normalization strategies and found that they exhibited a high correlation with each other, all having a Pearson correlation greater than 0.98 (Supp. Figure 5A). Due to the high correlation, we solely focused on the inferred binding weights using MatrixRider with a window frame of 1,000 bp upstream of the TSS for the comparison to the unweighted CollecTRI. We inferred TF activities utilizing the calculated binding weights and evaluated them in the benchmark framework as described above. Here, both the weighted and unweighted CollecTRI performed comparable (t-test: adjusted p-value equal to 0.87 and 0.94 for AUROC and AUPRC, respectively) (Supp. Figure 5B). Subsequently, we assessed the impact of pruning edges with low weights from the network and compared that with randomly removing edges. Our analysis revealed that this approach did not improve the network's performance for both the AUROC and AUPRC values in all cases (t-test: adjusted p-value > 0.05) (Supp. Figure 5C). In total, the weighting of CollecTRI's TF-gene interactions did not contribute to the improvement of CollecTRI's performance and is so far only possible for a subset of TF-gene interactions.

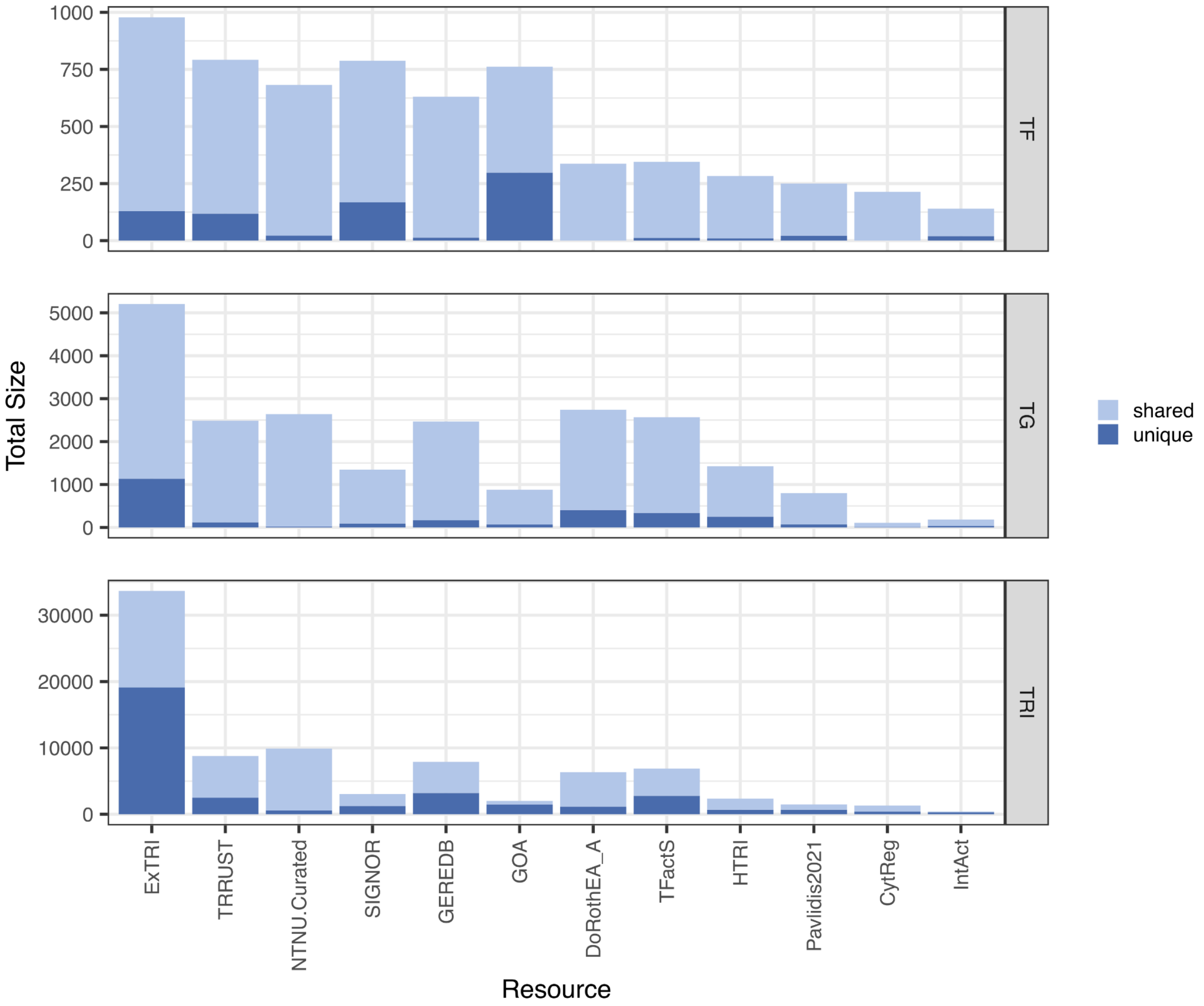


Supplementary Figure 1 ***Coverage of resources included in the CollecTRI meta-resource.*** *Number of unique and shared transcription factors (TFs) (top), target genes (TGs) (middle) and transcriptional regulatory interactions (bottom) and interactions (bottom) across the different resources which are part of CollecTRI. Any TF or interaction present in more than one resource is considered shared.*

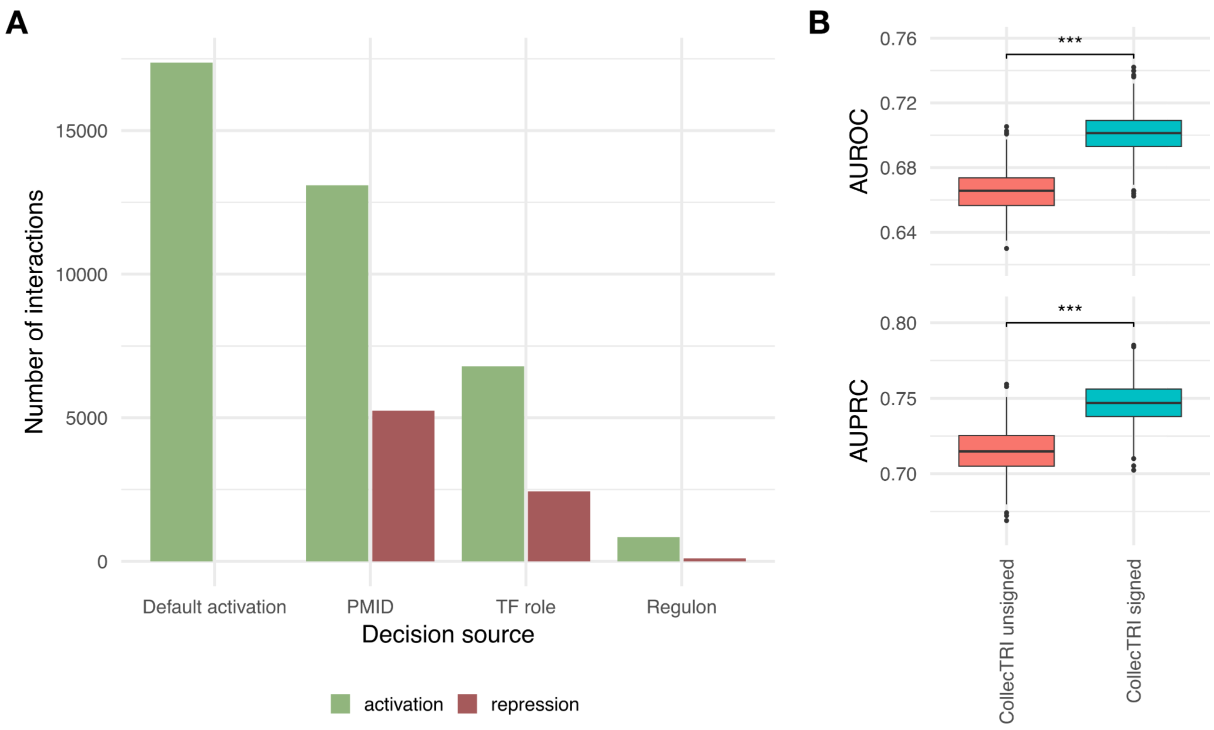


Supplementary Figure 2 ***Description of assigning the sign of regulation to the transcription factor (TF)-gene interactions.***

*(A) Overview of the number of interactions with an assigned mode of regulation based on different sources. The mode of regulation was assigned based on the prevalence of PubMed references (PMIDs), the general knowledge of a TF's role in activation (either activator or repressor), the mode of regulation of other genes in the TF's regulon or if no information was available an activating mode of regulation was assigned by default. (B) Predictive performance of the signed and unsigned CollecTRI-derived regulons. In the signed CollecTRI regulons a mode of regulation was assigned based on the sources described in (A). Predictive performance was measured by identifying perturbed TFs in knockTF experiments with AUROC (left) and AUPRC (right) for classifying TFs as perturbed or non-perturbed based on their estimated activities.*

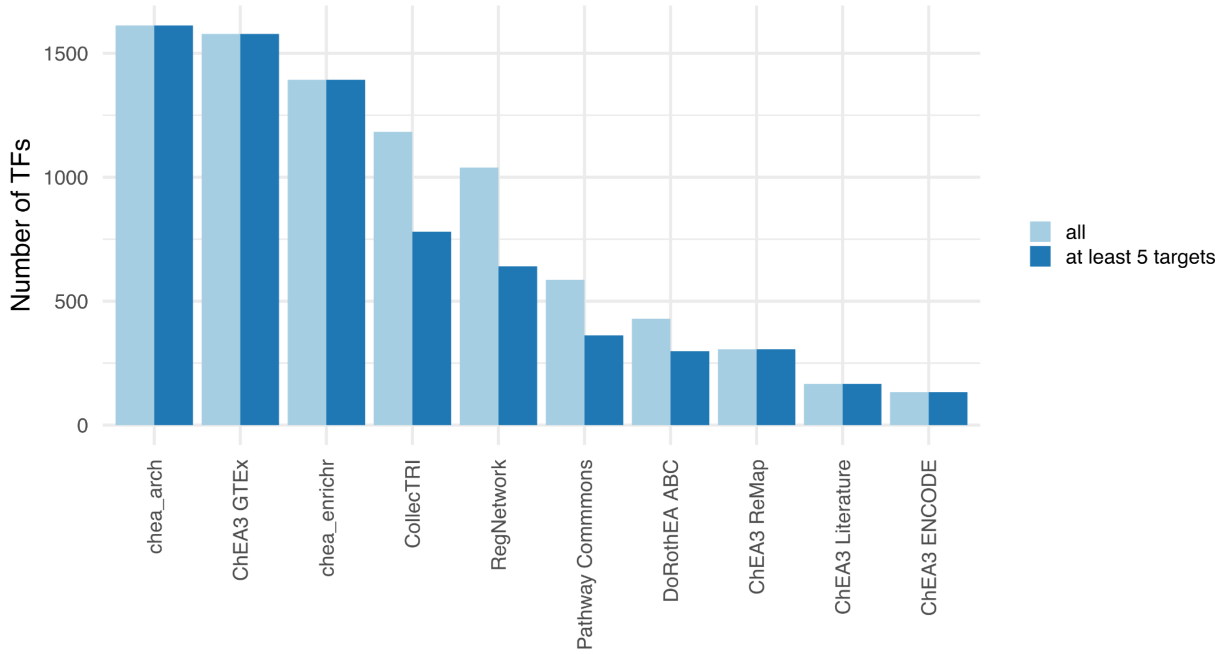


Supplementary Figure 3 ***Overview of transcription factors (TFs) across meta-resources.***
*Total number of TFs and number of TFs with at least five target genes present in each meta-resource.*

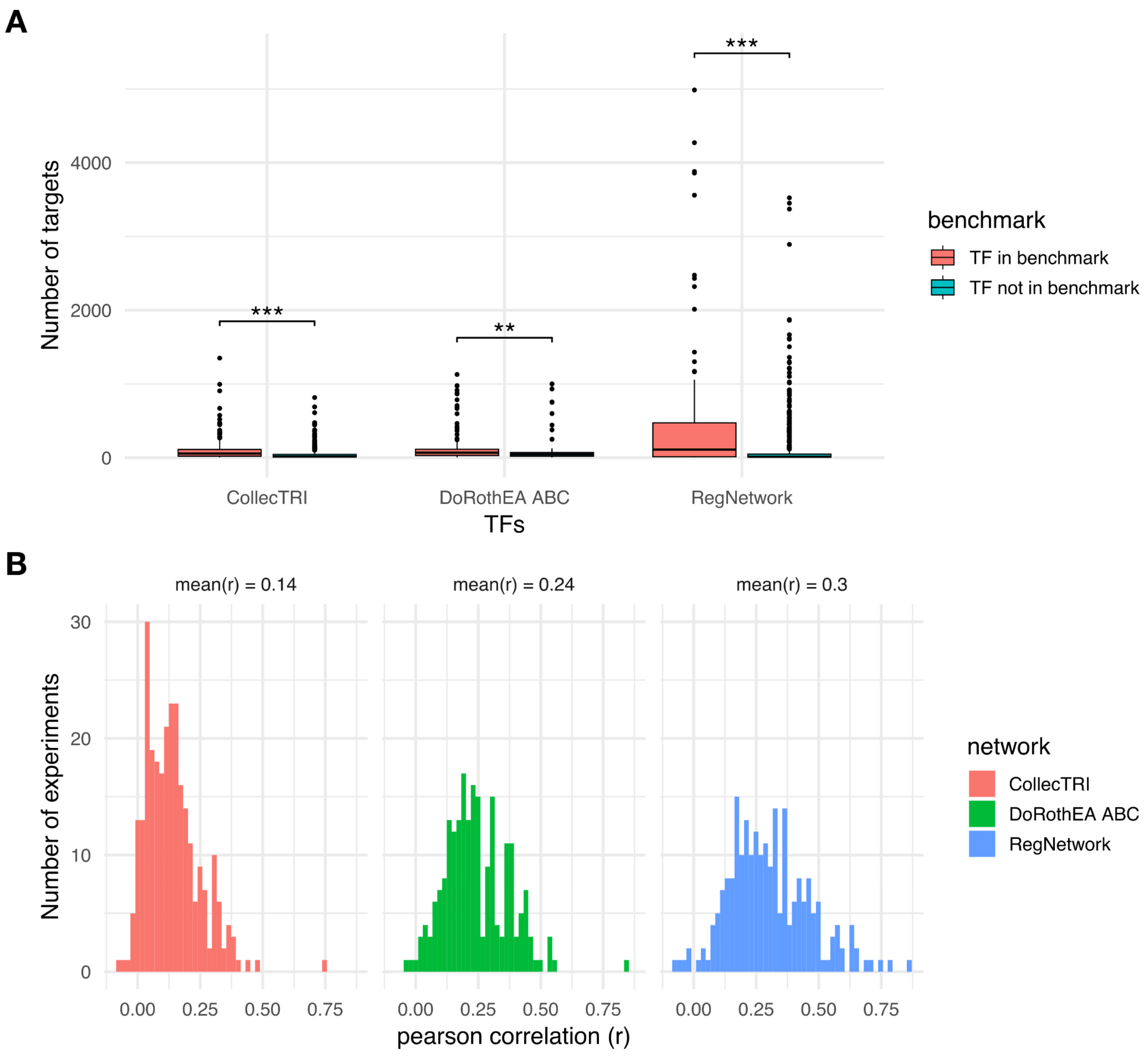


Supplementary Figure 4 ***Comparison between the number of targets of a transcription factor (TF) and its inferred activity.***

*(A) Comparison between the number of targets of TFs with and without available perturbation experiments in the benchmark. (B) Pearson Correlation between the number of targets and the activity of a TF for each experiment.*

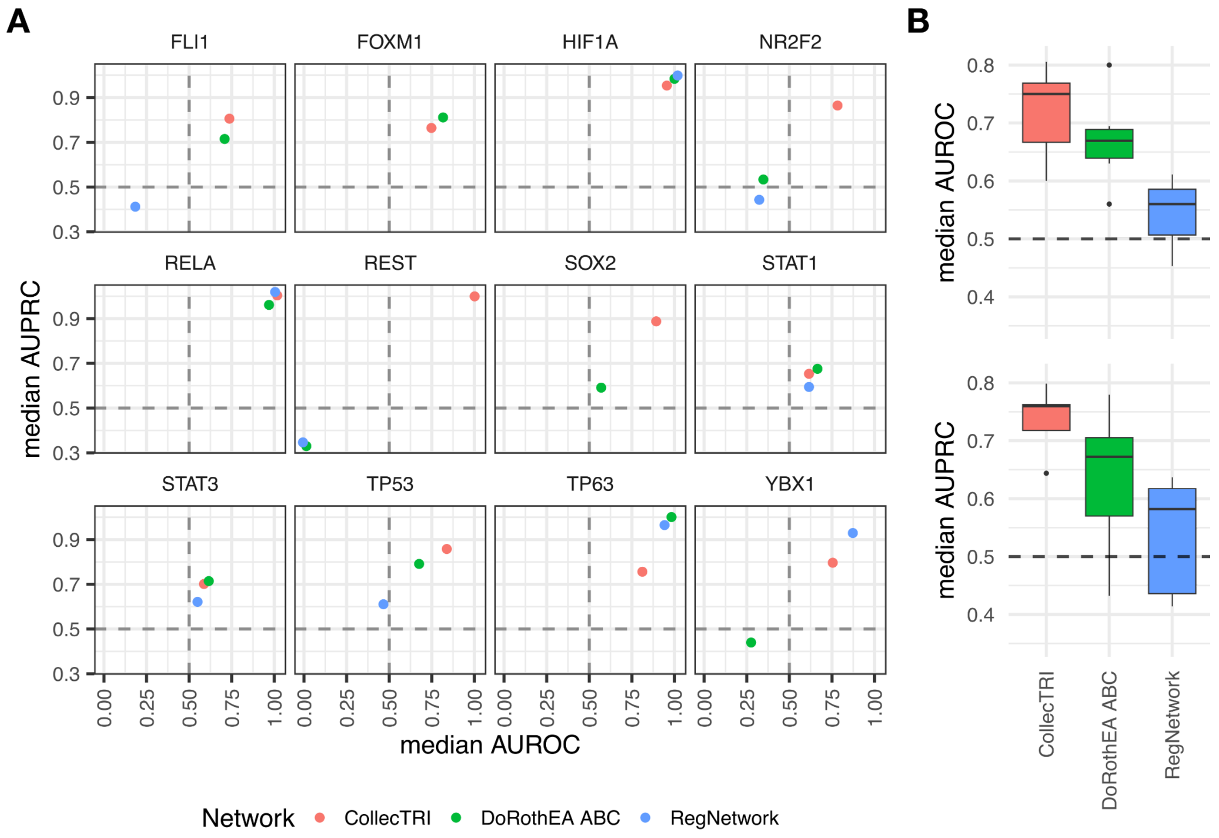


Supplementary Figure 5 ***Predictive performance to identify perturbation experiments for specific transcription factors (TFs).***

*(A) Predictive performance of regulon collections identifying perturbation experiments for a given TF. Only TFs with at least five perturbation experiments were selected. Median AUROC and median AUPRC for each regulon collection and each given TF classifying experiments as perturbed or non-perturbed. (B) Median AUROC (top) and AUPRC (bottom) across experiments for each regulon collection.*

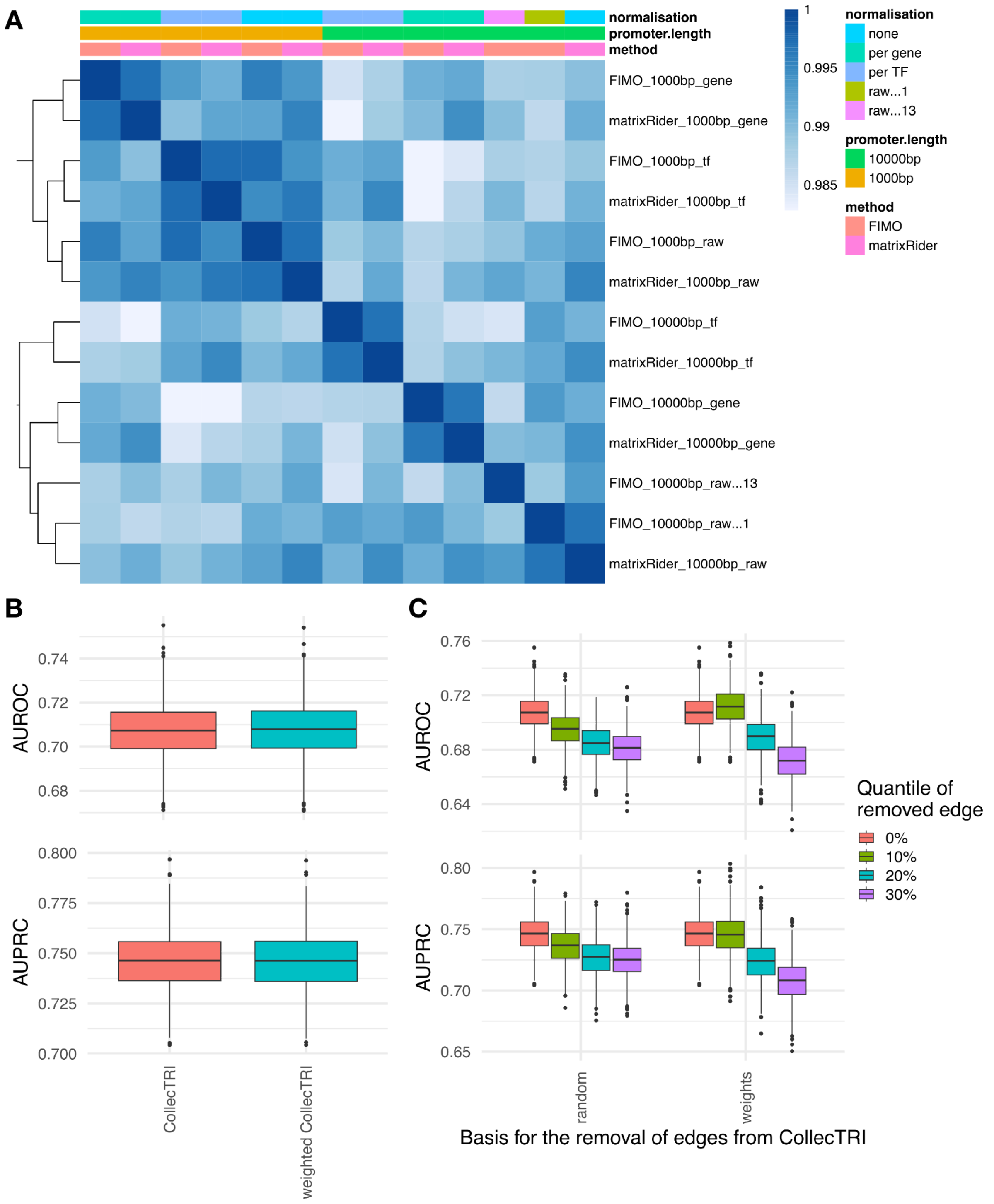


Supplementary Figure 6 ***Investigating the impact of weighting transcription factor (TF) – gene interactions in CollecTRI.***

*(A) Pearson correlation between inferred binding weights across tools, window sizes and normalization strategies. The tools MatrixRider and FIMO were used to calculate binding weights for a window size of 1,000 bp and 10,000 bp downstream of the transcription start site. The calculated weights were then normalized per gene or per TF and the final weights for each TF-gene interaction compared with Pearson Correlation. (B) Predictive performance of weighted and unweighted CollecTRI. Raw weights calculated from MatrixRider for a window frame of 1,000 bp were used for the comparison. (C)*

*Predictive performance of CollecTRI regulons after filtering TF-gene interactions with a low weight. Weights were prepared as described in (B) and the removal of the lowest 10%, 20% and 30% of TF-gene interactions was compared to randomly removing the same amount of edges.*
